# Wnt/β-Catenin Signaling Controls Spatio-Temporal Elasticity Patterns in Extracellular Matrix during *Hydra* Morphogenesis

**DOI:** 10.1101/214718

**Authors:** Mariam Veschgini, Hendrik O. Petersen, Stefan Kaufmann, Wasim Abuillan, Ryo Suzuki, Manfred Burghammer, Suat Özbek, Thomas W. Holstein, Motomu Tanaka

## Abstract

Albeit ample evidence has suggested the remodeling of extracellular matrix (ECM) in animals plays crucial roles in development and diseases, little is understood how ECM mechanics correlates with tissue morphogenesis. In this study, we quantitatively determined how spatio-temporal elasticity patterns in ECM change during the asexual reproduction of freshwater polyp *Hydra*. We first determined the mesoscopic protein arrangement in *Hydra* ECM (mesoglea) by grazing-incidence small-angle X-ray scattering with nano-beam (nano-GISAXS). Our data unraveled fibrillar type I collagen in *Hydra* mesoglea (Hcol-I) takes an anisotropic, more strongly distorted hexagonal lattice compared to those in vertebrates that could be attributed to the lower proline content and lack of lysin-crosslinks in Hcol-1 fibers. Then, we “mapped” the spatio-temporal changes in ECM stiffness *ex vivo* with aid of nano-indentation. We identified three representative elasticity patterns during tissue growth along the oral-aboral body axis of the animals. Our complementary proteome analysis demonstrated that the elasticity patterns of the ECM correlate with a gradient like distribution of proteases. Perturbations of the oral Wnt/β-catenin signaling center further indicated that ECM elasticity patterns are governed by Wnt/β-catenin signaling. The *ex vivo* biomechanical phenotyping of *Hydra* mesoglea established in this study will help us gain comprehensive insights into the spatio-temporal coordination of biochemical and biomechanical cues in tissue morphogenesis *in vivo*.

## Introduction

The extracellular matrix (ECM) regulates the homeostasis of animal tissue, supporting the structural integrity and cell functions [1]. ECM remodeling plays vital roles in tissue morphogenesis because it is accompanied by large-scale deformations, such as invagination, immigration and convergent extension during gastrulation [2–4]. Many diseases are also characterized by the significant remodeling of ECM, such as the myelofibrosis in bone marrow causing pancytopenia [5] and stiffening of pulmonary ECM in fibrotic lung tissues [6]. However, despite of accumulating knowledge on participating proteins and key signaling pathways, little is understood how ECM stiffness correlates with tissue morphogenesis [2]. Barriga et al indented cut pieces of *Xenopous laevis’* mesoderm with atomic force microscopy (AFM) and demonstrated by using head mesoderm of different age as well as hydrogels that the collective migration of neural crest cells during morphogenesis requires a stiffening of mesoderm [7, 8]. However, direct stiffness measurements of the ECM as a function of morphogenesis have not been reported so far. The use of the model animal with a simpler body design is therefore a straightforward strategy.

In this paper, we investigated the time progression of the ECM stiffness of fresh water polyp *Hydra* during tissue growth. *Hydra* is a member of a > 600 million years old phylum Cnidaria and a paradigm for an almost unlimited growth and regeneration capability. Compared to ‘higher’ animals, it has a simple, sack-like body plan with a body wall composed of an ECM, called mesoglea, which separates two cell layers; an outer ectoderm and an inner endoderm. Previous accounts showed that the mesoglea plays an important role in asexual reproduction of *Hydra* through budding. Mesoglea undergoes dynamic remodeling in order to support the daughter animal (bud) stemming out of the main body axis of a mother *Hydra* [9–11].

Intriguingly, the molecular composition of *Hydra* mesoglea is very similar to those of vertebrate ECM, containing heparan sulfate, laminin and fibronectin-like molecules as well as fibrillar (type I and II) and non-fibrillar (type IV) collagens [11–13]. *Hydra* ECM combines two major ECM functions that are separated in higher animals, i.e. mechanical integrity guaranteed by collagen fibers in connective tissues, and cell-cell communications mediated by basal lamina [13]. To accommodate both functions, *Hydra* mesoglea was supposed to form a tri-laminar structure: two thin sub-epithelial zones based on collagen type IV and laminin sandwich a central fibrous zone, the interstitial matrix consisting of a grid of collagen type I fibrils (Hcol-I) [14, 15]. However, despite the similarity in molecular compositions, there has been no quantitative comparison of the supramolecular organization between *Hydra* and vertebrates. Thus, we performed grazing-incidence small-angle X-ray scattering of native *Hydra* mesoglea by illuminating the mesoglea with a nano-focused beam (nano-GISAXS) [16, 17]. The use of a nano-beam (diameter: 200 nm) realized a much smaller beam footprint than a mesoglea, which enables one to gain the mesoscopic arrangement of ECM proteins in *Hydra* mesoglea in the directions parallel and perpendicular to the oral-aboral body axis (OA-axis). By scanning the mesoglea along the OA-axis, we unraveled that the tightly packed, fibrillar proteins form a distorted hexagonal lattice, whose unit lengths are distinctly longer to those in vertebrates. Moreover, the scattering signals collected from the direction perpendicular to the OA body axis suggested a clear anisotropy of fibrillar ECM proteins in *Hydra* mesoglea withstanding the significantly asymmetric stretching of *Hydra* body along the OA axis.

The dynamic morphological change during budding is supported by the remodeling of ECM by expression of proteases [18, 19]. Immunofluorescence labeling of type I collagen actually implied that the mesoglea of evaginating bud showed much weaker signals compared to the mother polyp. This can be attributed to the thinning of mesoglea in order to accommodate a large number of cells flowing out from the parental *Hydra* [20]. The thinning of ECM naturally suggests that rapidly growing buds are softer, but there has been no quantitative study on how tissue morphogenesis of *Hydra* correlates the local ECM stiffness during asexual reproduction. To extract characteristic spatio-temporal patterns of ECM stiffness during the morphogenesis, we isolated *Hydra* mesoglea at different time points from a synchronized culture and measured the local stiffness of *Hydra* mesoglea along the body axis by atomic force microscopy (AFM) nano-indentation. Different to whole animal explants [8], where cut pieces of tissue are always a function of the ECM and the overlying cells, this approach allows direct biophysical measurement of the ECM. The time progression of elasticity patterns along the body axis suggested that the break of the radial body symmetry in *Hydra* during budding is accompanied by a coordinated softening of the upper gastric region and stiffening of the budding region. The complementary proteomic analysis implied a significant change in protease expression patterns along the body axis. Time progression of the elasticity patterns from transgenic *Hydra* overexpressing β-catenin as well as wildtype *Hydra* treated with GSK3β inhibitor (Alsterpaullone) demonstrated the reduced stiffness of *Hydra* ECM correlates with the activation of canonical Wnt signaling.

## Results

### Mesoscopic arrangement of *Hydra’s* ECM

In a first approach we determined the mesoscopic order of *Hydra* ECM (mesoglea) by using nano-GISAXS. Figure 1a schematically illustrates the experimental setup and sample geometry. The *Hydra* mesoglea isolated by freeze-thawing (see Materials and Methods) was deposited on a Si3N4 membrane. A monochromatic, nano-focused beam (beam diameter: 200 nm) impinges to the sample at an incident angle *α_i_* = 0.46°, which is beyond the critical angle of total external reflection (*α_c_* = 0.14°). Figures 1b and 1c represent nano-GISAXS patterns obtained from the directions parallel and perpendicular to the body axis of mesoglea, respectively.

**Figure 1.**
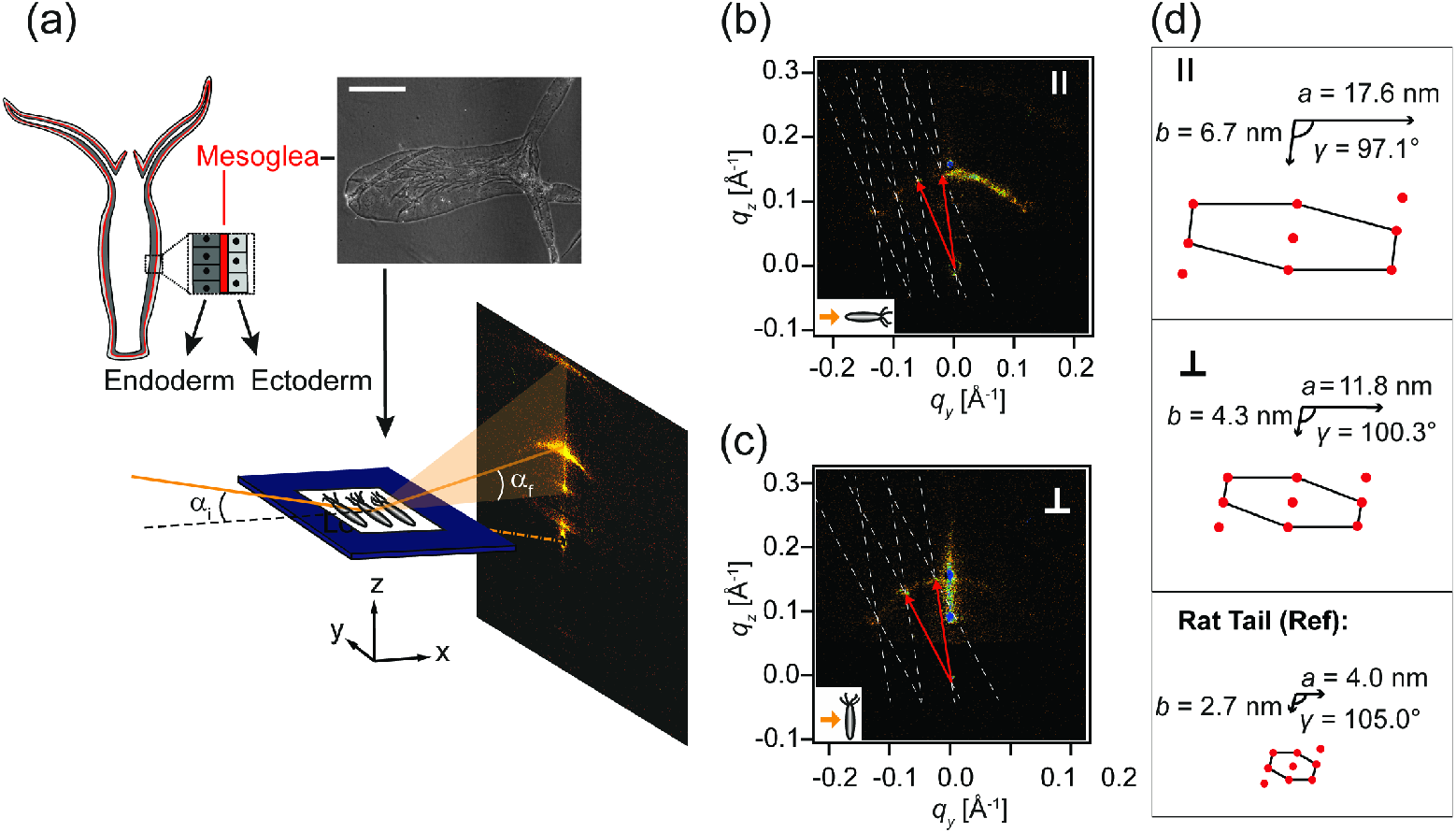
Nano-GISAXS of *Hydra* ECM mesoglea. (a) Top left: body design of sweet water polyp *Hydra*. Mesoglea (red) was isolated by freeze thawing. Top right: phase contrast microscopy image of an isolated mesoglea. Scale bar: 500 μm. Bottom: experimental setup of nano-GISAXS. Isolated mesoglea was placed on a Si3N4 window and illuminated by a nano-focused X-ray beam (diameter: 200 nm) at a grazing incidence angle *α_i_* = 0.46°. Scattering patterns obtained from mesoglea, whose body axis was positioned parallel (b) and perpendicular (c) to the beam (see inset yellow arrow). The reciprocal lattice is indicated in white, and the lattice vectors in red. (d) Lattice parameters in real space (*a, b, γ*) calculated from the directions parallel (top) and perpendicular (middle) to the major body axis. For comparison, the lattice parameters from the reference sample (collagen type I from rat tail tendon) [21] are presented (bottom).

Several scattering patterns showed that one or more arcs were arranged concentrically around the direct beam, accompanied by additional satellite peaks positioned on the arcs (Figure1d). The white grid overlaid on each pattern represents the reconstructed reciprocal lattice. The obtained lattice patterns clearly indicate that rod-like objects are aligned parallel to the substrate surface. The real space lattice parameters (*a, b, γ*) calculated from the GISAXS patterns parallel and perpendicular to the *Hydra* body axis are presented in the top and middle panels of Figure 1d, respectively. The lattice parameters of the mesoglea parallel to the OA-axis are *a*_║_ = (17.6 ± 3.6) nm, *b*_║_ = (6.7 ± 0.8) nm and *γ*_║_ = (97.1 ± 2.4)°, implying that the rods take a distorted hexagonal arrangement. The corresponding data show that the rod-like objects also take an ordered structure perpendicular to the body axis resulting in a distorted hexagonal lattice with slightly different units, *a*_⊥_ = (11.8 ± 0.5) nm, *b*_⊥_ = (4.3 ± 0.4) nm and *y*_⊥_ = (100.3 ± 9.3)°. We compared these lattice parameters with lattice parameters of type I collagen from rat tail tendon taken from a previous account (*a* = 4.0 nm, *b* = 2.3 nm and *γ* = 105.0° [21], bottom panel of Figure 1d). Similar opening angles (≈ 100 °) suggest that the rod-like objects in *Hydra’s* mesoglea are fibrillar type I collagen (Hcol-I), which is the main component of the middle layer of *Hydra’s* mesoglea [14, 15]. Clearly different lattice parameters between the directions pararell and perpendicular to the OA axis indicate that the mesoscopic arrangement of Hcol-I fibers is highly anisotropic, which enables Hcol-I to support the large degree of anisotropic extension and contraction along the body axis. Moreover, the interfibrillar distances between Hcol-I are about 2 – 4 times longer compared to those in rat tail tendon. This can be attributed to the lower proline content and lack of lysin-crosslinks in Hcol-I fibers [15, 22]. Thus, our data on the mesoscopic structural order of *Hydra’s* mesoglea clearly indicate a dominant role of fibrillar type I collagen (Hcol-I) in the generation of the basal stiffness of *Hydra’s* mesoglea.

### Elasticity mapping and classification of mechanical phenotypes

To monitor the change in mesoglea stiffness during the asexual budding, we mapped the stiffness (bulk elastic modulus) of *Hydra* mesoglea along the body axis by AFM nano-indentation [23–25]. Figure 2a shows a typical force-distance curve of *Hydra* mesoglea. Note that the readout from the instrument presented in Y-axis is given in voltage, which is proportional to the force. We fitted the measured data (grey symbols) with the modified Hertz model [26] and calculated the bulk elastic modulus. Prior to the analysis, we carefully optimized the tip-sample contact point *zo*, as the calculated elastic modulus significantly depends on the accuracy of *z*_0_ [27, 28]. As indicated by arrows in Figure 2a, the contact point candidate was marched over the points near the onset point to determine the optimal contact point (red). As presented in Figure 2b, we determined the elastic modulus *E* by minimizing the mean sum of square residuals (SSR). For each sample, 7 – 37 positions were indented. As shown in Figure 2c, the elasticity pattern was extracted by normalizing each indentation point with the relative position from the foot (*d* = 0) to mouth (*d* = 1.0) along the body axis. This enables one not only to compare different mesoglea samples but also to discern three key regions along body axis: (i) peduncle region (0.0 ≤ *d* ≤ 0.1), (ii) budding region (0.1 ≤ *d* ≤ 0.3), and (iii) upper gastric region (0.6 ≤ *d* ≤ 1.0).

**Figure 2.**
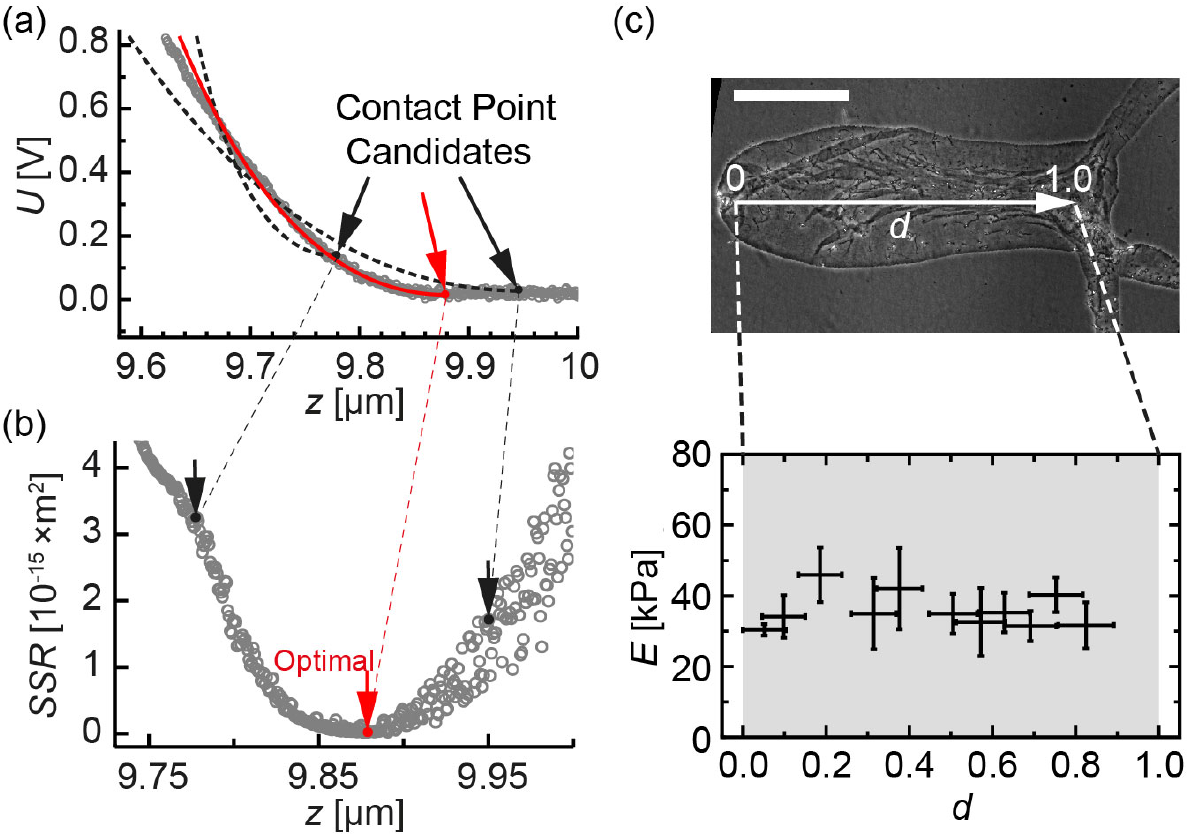
Elasticity mapping of *Hydra* mesoglea. (a) A typical force (voltage) – distance curve (gray circles) of a mesoglea isolated from a freshly detached *Hydra*. Three contact point candidates (indicated by arrows) and the corresponding fits are presented for comparison. (b) The optimization of the fit by minimizing the mean sum of square residuals (SSR) during the marching of the contact point candidate. The optimal contact point (red) yields the bulk elastic modulus *E* = 29.7 kPa. (c) Phase contrast microscopy image of a mesoglea isolated from a freshly detached *Hydra*. The relative position from the foot (*d* = 0) to mouth (*d* = 1.0) along the body axis is used for the normalization of spatial information. Scale bar: 500 μm. (d) “Elasticity map” of *Hydra* mesoglea along the body axis. Error bars correspond to the standard deviation out of 8 independent measurements.

Utilizing the positional segmentation described above, the “elasticity maps” extracted from different animals could be classified into three major elasticity patterns (Figure S1): ***Type A*** (Figure 3a) is characterized by uniform elastic moduli along the entire body column with *E* ≈ 40 kPa. ***Type B*** (Figure 3b) exhibits distinctly higher elastic moduli in the peduncle and budding regions (*E* ≈ 120 kPa) compared to the head region, *E*≈ 20 kPa. ***Type C*** (Figure 3c) looks similar to type B but shows lower elastic moduli in the peduncle region, *E* ≈ 50 kPa. The overlay of the three main elasticity patterns is presented in Figure 3d. 97 % of the measured elasticity patterns in mesoglea isolated from *Hydra* animals (*n* = 38) at different time points could be classified into type A (66 %), type B (18 %), and type C (13 %). Only one sample exhibited a random pattern, which could be attributed to the artifacts from remaining nematocysts (data not shown).

**Figure 3.**
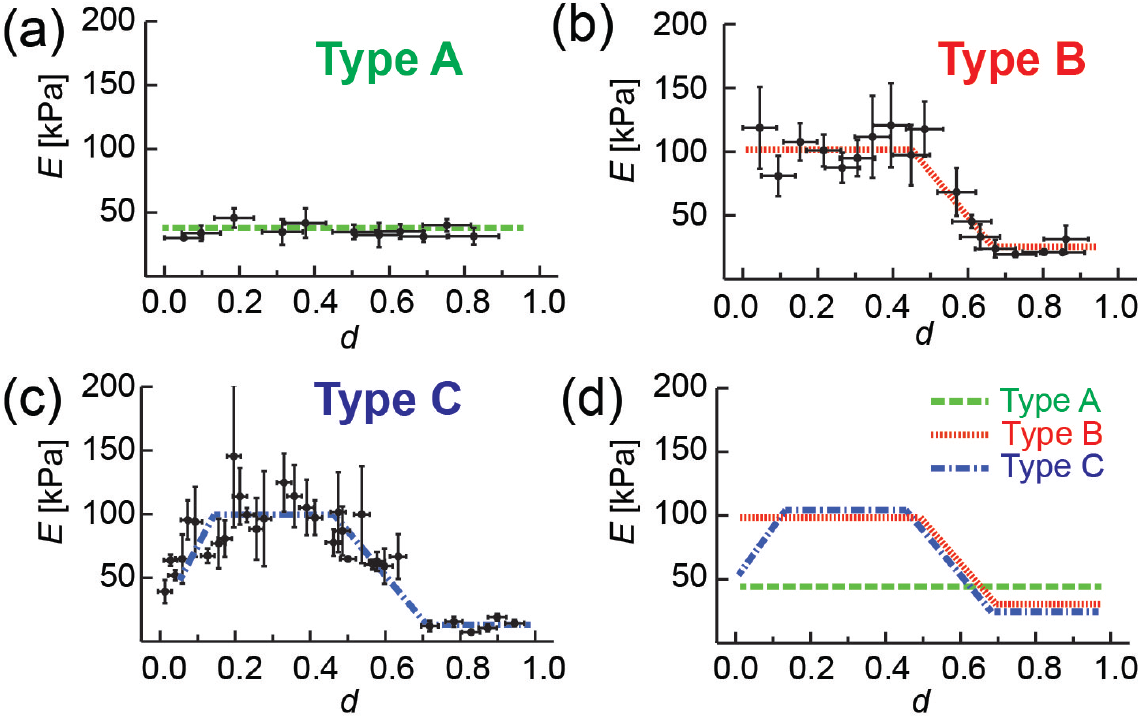
Three representative elasticity phenotypes of *Hydra* mesoglea (*n* = 38). (a) Type A is characterized by a uniform, soft mesoglea. (b) Type B shows higher elastic moduli in peduncle and budding regions compared to the head region. (d) Type C looks similar to type B, but the elasticity in the peduncle region is distinctly lower. (d) Overlay of three representative elasticity patterns.

### Modulation of elasticity patterns correlates with ECM remodeling

To unravel whether the extracted elasticity patterns correlate with the age or developmental stage of *Hydra*, we compared the elasticity patterns of freshly detached animals (*t* = 0 d) and animals at the ages of *t* = 2 - 3 d. The time window of 3 d was chosen in order to focus on the transition from newly detached buds to budding animals. Mesoglea from freshly detached *Hydra* animals (*t* = 0 d) showed exclusively type A pattern, i.e. the elasticity of mesoglea was low and uniform along the body axis. At *t* = 2 d, mesoglea started taking type B and type C patterns (≈ 60 %), indicating the increase in the ECM elasticity in the budding region. At *t* = 3 d, type B and type C were predominant (≈ 80 %). Although it is difficult to identify the very early stage of bud formation, the emergence of type B and type C can be attributed to the formation of buds. In fact, *t* = 2 – 3 d is the typical time window for freshly detached *Hydra* animals to start budding [29]. ECMs of fast-growing buds became thinner by time [20], resulting in a homogeneous and low elasticity pattern (type A). Our data indicate that the growth process and maturation of freshly detached buds towards mature, bud-forming animals is accompanied by the modulation of elasticity patterns in *Hydra* ECM. It seems plausible that the growth of buds from mature *Hydra* animals is supported by the stiffening of ECM in the budding region, which is characterized by the conversion from type A to type B/C. Last but not least, the low stiffness levels in type C at the aboral end likely correlate with the maturation of polyps undergoing dynamic remodeling of peduncle region [30].

The molecular processes that enable the large-scale deformation must include either post-transcriptional modifications of ECM proteins or changes in expression and composition of ECM proteins (Figure 4a). To unravel the molecular-level regulation of mesoglea elasticity both in space and in time, we performed a quantitative proteomic analysis of *Hydra* mesoglea. We compared the protein expression patterns of the mesoglea from freshly detached buds and mature *Hydra* animals. For mesoglea from freshly attached animals (*t* = 0 d), we analyzed whole animals due to the limited sample availability. On the other hand, for mesoglea from older and budding animals (*t* = 3 d), we separated the samples into two groups; ECM from the upper gastric and head region, and ECM from the lower gastric and budding region. As the internal standards, we used isolated mesoglea samples from SILAC *Hydra* containing all developmental stages [31]. Firstly, we found that the core members of the mesoglea proteins [13] showed no detectable changes between the three sample groups. This finding indicates that the remodeling of ECM proteins but not the change in ECM composition is responsible for the modulation of elasticity patterns. However, we found a significant down-regulation of four proteins in the mesoglea from the lower gastric and budding region of older and budding polyps: Dickkopf-1/2/4 protein (Dkk 1/2/4) [32, 33], a GM2 activating protein [34, 35], and two matrix proteases, i.e. metallo- and Astacin-like proteases. From a functional viewpoint, two matrix proteases are potential candidates for this mechanism (Figure 4b). Both proteases were strongly abundant in the mesoglea of upper gastric and head region from 3 d-old polyps and freshly detached buds (grey). These data are fully consistent with the elasticity patterns showing (i) a lower stiffness of the mesoglea from freshly detached polyps and from the upper gastric region and (ii) a much higher stiffness of the mesoglea from the lower gastric and budding region. This provided direct evidence that the modulation of elasticity patterns in *Hydra* mesoglea are driven by the expression patterns of identified matrix proteases.

**Figure 4.**
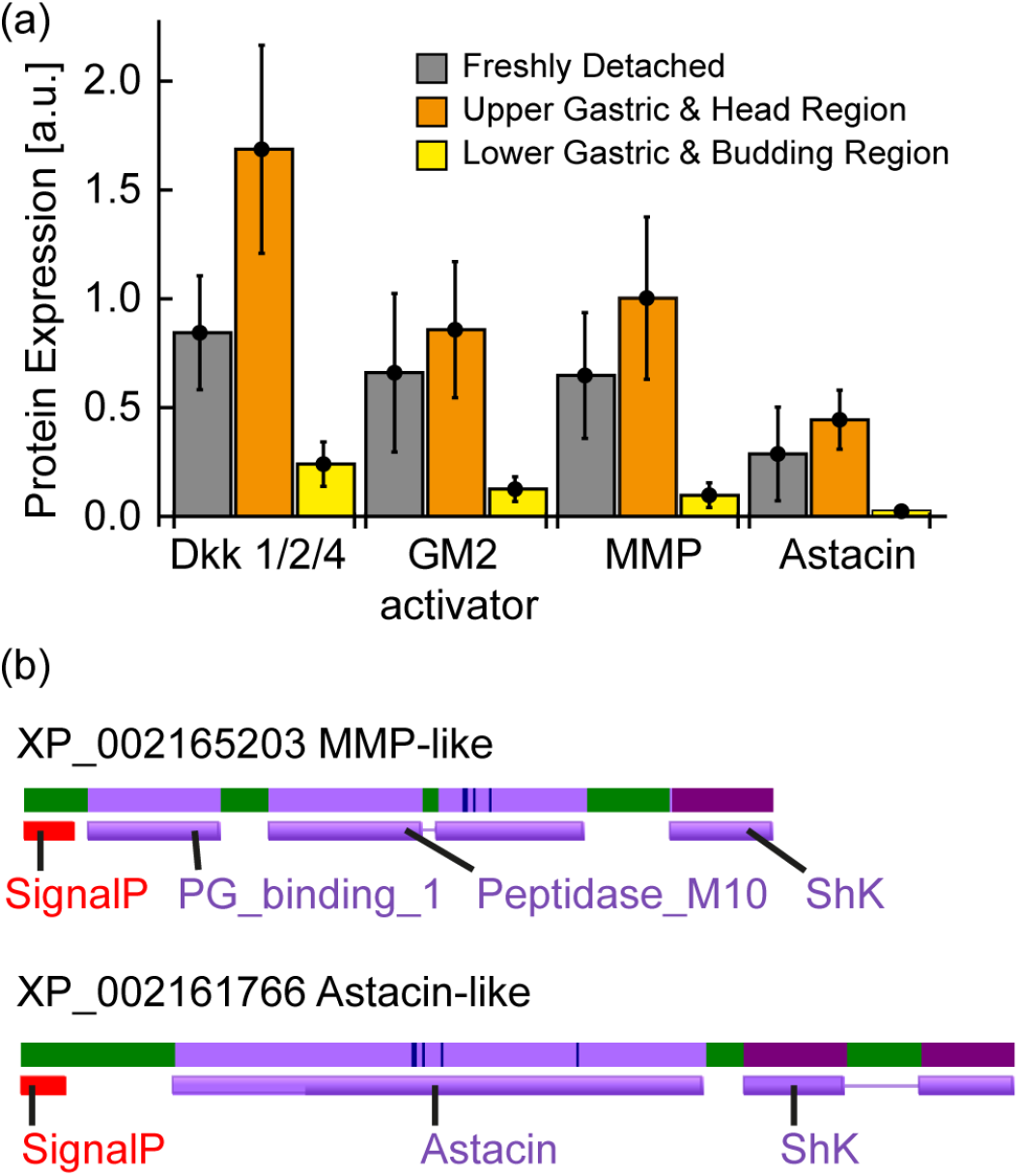
Mesoglea proteome analysis. (a) Protein expression difference was analyzed between mesoglea of freshly detached animals (*t* = 0 d, grey) and mesoglea from mature animals. The mesoglea from mature animals (*t* = 3 d) were separated into that form upper gastric and head region (orange) and lower gastric and budding region (yellow). Multiple sample ANNOVA analysis, using a p-value threshold of 0.05, revealed significant expression differences of four proteins between the samples. (b) Domain structures of MMP- and Astacin-like proteases.

### Elasticity patterns correlates with Wnt/β-catenin signaling

Notably Dkk-1/2/4 is expressed in a gradient-like pattern along the oral-aboral-axis of *Hydra* and acts as an antagonist against Wnt ligands [32, 33]. We therefore analyzed to what extent the perturbation of the Wnt/β-catenin pathway can modulate the elasticity patterns of *Hydra’s* mesoglea. *Hydra* animals were treated with Alsterpaullone (Alp) [36]. Alp is an inhibitor of glycogen synthase kinase 3β (GSK3β), inducing the stable activation of β-catenin in a position-independent manner [37, 38]. The treatment of *Hydra* animals with 5 μM Alp leads to the formation of ectopic tentacles and head structures along the body column after 2 - 3 d [36]. Figure 5a shows an optical microscopy image of an Alp-treated *Hydra* polyp (see Materials & Methods). Compared to untreated controls, the mesoglea of Alp-treated animals exhibited a dramatic decrease in its stiffness along the entire body column to *E* ≈ 35 kPa (Figure 5b). We also determined the elasticity patterns in mesoglea isolated from β-catenin overexpressing *Hydra vulgaris* that form ectopic tentacles and head structures [39]. As presented in Figure S2, the mesoglea of β-catenin overexpressing *Hydra* exhibited the same drop, although the elastic moduli were more diverging due to a broader diversity of morphological phenotypes and heterogeneous expression levels of β-catenin in individual animals. Our data show that there are unexpected changes in the elasticity patterns of *Hydra’s* mesoglea during *Hydra* morphogenesis that are tightly controlled by Wnt/β-catenin signaling. The elasticity patterns in *ex vivo* ECM unraveled in this study will help us understand how biochemical and biophysical cues are managed in space and in time during tissue morphogenesis *in vivo*.

**Figure 5.**
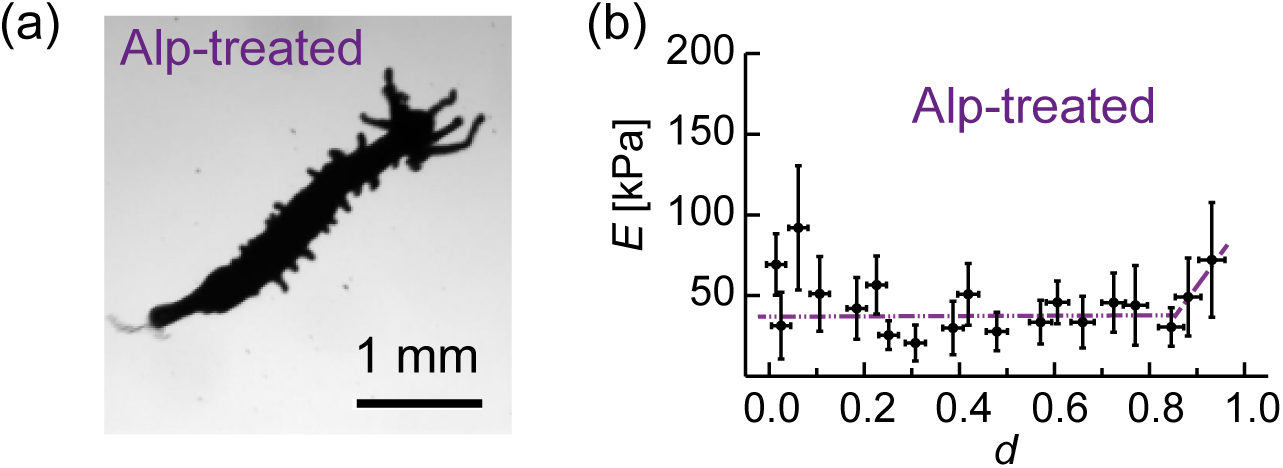
(a) Light microscopy image of *Hydra magnipapillata* treated with GSK3β inhibitor, Alsterpaullone (Alp) at *t* = 3 d. (b) The same animal treated with Alp at *t* = 3 d. (c) Elasticity pattern of mesoglea isolated from Alp-treated *Hydra*. The elastic moduli were low over the entire body column, which is clearly different from intact mesoglea (Figure 3).

## Discussion

The ECM in multicellular organisms serves as a cell substrate and as a medium for the transport of extracellular factors required for coordinated growth and patterning. Mounting evidence suggested that the remodeling of ECM, such as mesoscopic structural arrangement and modulation of ECM stiffness, is one of the prominent features in development and diseases [1]. In this study, we focused on how the patterns of ECM elasticity are spatio-temporally modulated in animals undergoing dynamic tissue morphogenesis. As the animal model, we selected fresh water polyp *Hydra*, which is a paradigm for an almost unlimited regeneration capability [31].

Firstly, we determined the mesoscopic arrangement of ECM proteins using nano-GISAXS, which has been used mainly for colloids and polymers [16, 17]. As nano-GISAXS enables one to collect the structural information from the beam footprint smaller than a mesoglea, we unraveled that collagen fibers (Hcol-I) take highly asymmetric, distorted hexagonal lattices. The packing of Hcol-I fibers along the body axis was found to be less ordered compared to the one perpendicular to the body axis, which can be attributed to an intrinsic body design of *Hydra* undergoing significant stretching and contraction along the oral-aboral axis. It is notable that the unit cell size of Hcol-I fibers is about 2 – 4 times larger compared to the standard samples from collagen type I fibers from rat tail tendon [21, 40, 41]. The apparently different packing of collagen fibers from vertebrates can be attributed to the amino acid composition. Compared to type I collagen from vertebrates, Hcol-I has a lower proline content and lacks typical lysine-crosslinking sites [15]. Moreover, an altered post-translational processing leads to the retention of the N-terminal pro-peptide-like domains [15], resulting in weaker interfibrillar interactions between Hcol-I. It is remarkable that the collagen of *Hydra’s* ECM actually shares common structural features with as Ehlers-Danlos syndrome characterized by the hypermobility of joints (which is claimed as the reason for extraordinary performance of Niccolo Paganini) [42].

Previously, Burnett and Hausmann reported that *Hydra* mesoglea can be stretched even up to 7 – 8 times its length and twice its width, and it recovers the normal shape upon releasing the stress [43]. Aufschnaiter et al. showed that *Hydra* mesoglea is continuously remodeled and moves towards both ends of the body column [20]. This displacement occurs largely in continuity with the epithelial cells that are secreting the ECM and is in accord with previous cell labeling data from Campbell [30, 44]. In the budding region, the parental mesoglea is recruited and stretched along evaginating buds without biosynthesis of new collagens at the early stages [20], suggesting a lower ECM stiffness in budding region. However, there have been no studies how the ECM remodeling affects the biophysical properties of the ECM in space and in time during large morphological changes. By quantitative comparison of the elasticity patterns of mesoglea samples isolated from the synchronized culture by AFM nano-indentation, we unraveled the emergence of three characteristic “mechanical phenotypes” of *Hydra’s* mesoglea during tissue morphogenesis. After detachment from the parental animal, the elastic moduli of the mesoglea from young polyps were low and uniform over the body axis (type A). The elasticity patterns of the mesoglea changed over 2 - 3 d, showing an increase in elastic moduli in budding region and a decrease in the gastric region (types B/C). This emergence of elasticity gradients from an initially uniform elasticity distribution over time seems plausible from the viewpoint of mechanics, because the mature ECM of the budding region must mechanically support a rapidly growing bud possessing high local curvatures. On the other hand, from the viewpoint of protein biochemistry, the emergence of elasticity gradient clearly indicates either post-transcriptional modifications or changes in expression and composition of ECM proteins. We performed a quantitative proteomic analysis of *Hydra* mesoglea of freshly detached buds (*t* = 0 d) and mesoglea of mature *Hydra* animals (*t* = 3 d) isolated from the upper gastric and lower budding regions. Firstly, we found no change in protein compositions, confirming that the modulation of elasticity patterns is driven by the remodeling of existing ECM proteins. Among the proteins showing distinct changes in expression, we identified MMP- and Astacin-like protease as the candidates that modulate the ECM stiffness. In fact, these proteases are distinctly down-regulated in the lower gastric and budding region of polyps, while the expression levels remained high in the upper gastric region. As schematically presented in Figure 6a, freshly detached animals (*t* = 0 d) possess uniformly soft mesoglea (type A), associated with moderate and uniform protease expression level. On the other hand, in budding polyps (*t* = 2 – 3 d), the down-regulation of protease expression in the lower gastric region leads to the stiffening of ECM, resulting in type B/C (Figure 6b). The softening of ECM in the peduncle region (type C pattern) can be explained by an up-regulation of metalloprotease near the foot [9] or bud. These results provide direct evidence that the modulation of elasticity patterns correlate with the remodeling of mesoglea by proteases.

**Figure 6.**
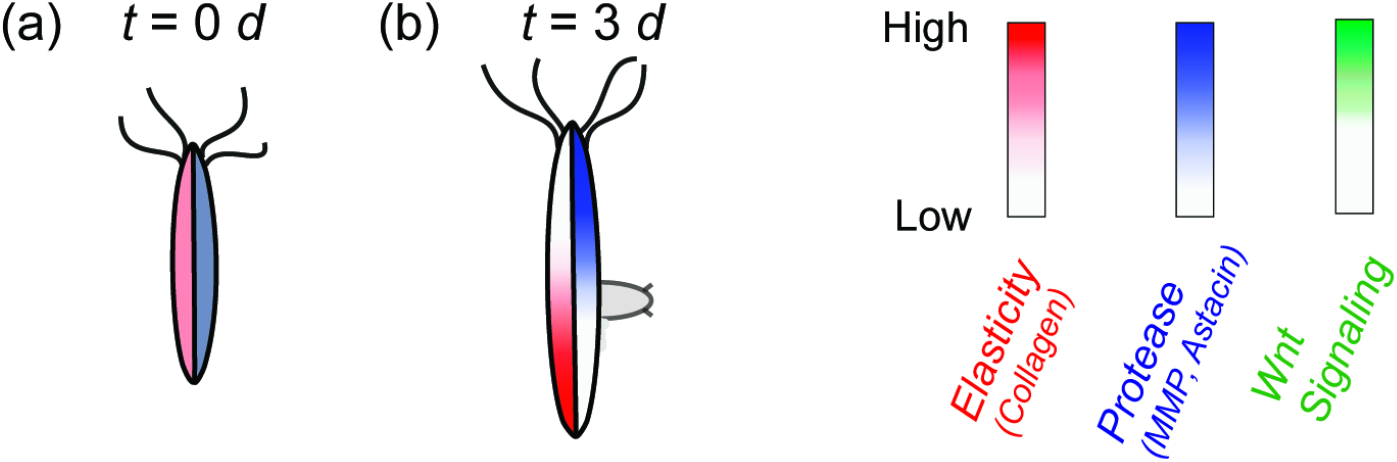
Spatiotemporal patterns of ECM elasticity and protease expression. (a) Freshly detached *Hydra* (*t* = 0 d) possesses uniform, soft mesoglea (type A), associated with moderate and uniform protease expression level. (b) According to aging (*t* = 3 d), the down-regulation in the budding region leads to the elevation of elastic moduli, resulting in type B/C patterns.

It has been established that Wnt/β-catenin signaling plays central roles in axis formation of *Hydra*. Other studies have also suggested that Wnt signaling induces the up-regulation of metalloproteases in various developmental processes [45–49], e.g. in embryonic neuronal stem cells [50], in human mesenchymal stem cells, but also in the migration of single T-cells [46] or in the collective migration of epithelial cells during would healing [45]. Therefore, we investigated whether the perturbation on Wnt/β-catenin pathway could influence the elasticity patterns in *Hydra* mesoglea by (a) treating wildtype polyps with Alsterpaullone (Alp, Figure 6) and (b) using β-catenin overexpressing *Hydra* animals (Figure S2). Alp is an inhibitor of GSKβ3, activating β-catenin all over the body column. Both Alp-treated and β-catenin overexpressing *Hydra* polyps exhibited significantly low elastic moduli all over the body column at *t* = 3 d. This is clearly different from the intact, wildtype *Hydra* exhibiting high elastic moduli in the budding region (type B/C). These results indicate that the modulation of elasticity patterns in mesoglea is tightly coupled to Wnt/β-catenin signaling.

The next question one can address is whether the ECM stiffening coincides with the formation of highly curved structures, such as buds and tentacles. Several in vitro studies suggested that the collective cell migration, such as would healing, is faster on stiffer substrates [51, 52]. In case of budding *Hydra* polyps, the movement of cells into buds during the early budding stage is coupled to that of mesoglea, which is distinctly different from the normal migration caused by cell growth [20]. As presented in Figure S3, the stiffening of ECM is reflected to the increase in the actin filaments in the budding region. It is notable that such an unusual movement was also observed in tentacle-forming tissues [20]. Thus, one could hypothesize that bud and tentacle forming tissues might exhibit a similar elasticity level, if the increased stiffness of the mesoglea simply reflects the biomechanical constraints to support the evagination of highly curved structures. However, our Wnt perturbation results implied the opposite. The activation of Wnt pathway resulted in uniformly soft ECM along the entire body column indicating a high level of MMP protease activity (Figures 6 and S2). Thus, we concluded that there is no direct correlation between the ECM stiffening and evagination of highly curved tissues.

Intriguingly, the lower gastric region, which is also characterized as the region where stem cells show high activity and proliferation capacity, turns out to exhibit a highly dynamic landscape in terms of the biophysical properties of its ECM. Regions with a lower or medium stiffness (i.e. the upper gastric region and the growing bud) are those regions that have a continuous and high proliferative capacity [53]. Currently, it is unclear whether these dynamic changes in ECM stiffness are also coupled to the stem cell functions of interstitial stem cells located in the upper and lower gastric region [54, 55]. Further *in vitro* studies using substrates with tunable stiffness [56, 57], *in vivo* modulation of ECM stiffness [8] and a genome-wide RNA tomography [58] along the OA axis of *Hydra* would help us unravel if the elasticity patterns in ECM are coupled to stem cell activities.

## Materials and Methods

### Hydra culture

*Hydra magnipapillata* and β-catenin overexpressing *Hydra vulgaris* were cultured in modified *Hydra* medium [59] (1.0 mM CaCl_2_, 1.0 mM Tris-HCl, 1.0 mM NaHCO_3_, 0.1 mM KCl, 0.1 mM MgCl_2_, pH = 7.4) and kept at (18 ± 0.5)°C. The culture was synchronized. Animals were fed daily with *Artemia* and cleaned a few hours after feeding. SILAC *Hydra* were produced as described earlier [31].

### Isolation of mesoglea

*Hydra* animals were starved for 1 d and transferred to a 2 ml test tube with *Hydra* medium (1 *Hydra* per tube). *Hydra* medium was replaced by 1.5 ml of a 0.5 % w/v solution of N-lauroylsarcosine sodium salt in water. The sample was then immersed in liquid nitrogen for 10 min. Frozen *Hydra* samples were either used directly or stored at - 80 °C. To extract the mesoglea *Hydras* were transferred to distilled water after thawing and forced through a Pasteur pipette 20 to 40 times. The washing water was exchanged 3 times. Subsequently the isolated mesoglea was transported to a dry Petri dish with a drop of water and allowed to dry overnight at room temperature.

### Nano-GISAXS experiments

Nano-GISAXS experiments were performed at the European Synchrotron Radiation Facility (ESRF) beamline ID13 (Grenoble, France). Mesoglea samples were prepared 2 – 3 d before experiments and deposited on a 100 nm-thick Si3N4 window surrounded by a Si frame (SPI supplier, United States). Experiments were performed on dehydrated samples and at *T* = 20 °C and ambient pressure. Mesoglea samples were probed with nano-focused beam (diameter: 200 nm) with a wavelength of 0.81 Å (15.3 keV) and at an incident angle of 0.46°. Data was collected with a 2D detector (Maxipix, ESRF, France) [60]. Silver behenate (C_21_H_43_COOAg) was used for calibration.

### Nano-indentation experiments

Nano-indentation was performed utilizing a JPK-NanoWizard3 atomic force microscope (JPK, Germany) mounted on a Zeiss Axiovert 200 microscope (Zeiss, Germany). Silicon nitride Cantilevers with a nominal spring constant of 10 mN/m and tip-radius of 20 nm were used (MLCT- Probes purchased from Bruker, United States). Indentation experiments were carried out in 150 mM NaCl. The spring constant of each cantilever was determined prior to measurements using the thermal noise method. A total number of *n* = 38 *Hydra* animals out of a daily fed synchronized culture were examined. *Hydra* samples were between 0 and 10 d old and bore no more than 3 buds. Depending on the size, each mesoglea was indented at 7 – 37 different positions along the body column. Mesoglea was indented with a speed of 2 – 6 μm/s and up to a cantilever deflection of 0.8 V relative to the baseline. At each position along the *Hydra* body axis 4 points with a distance of 5 μm were indented.

Each spot was indented two times. Subsequent measurements at the same spot with a standard deviation larger than 10 % were discarded. Local elastic moduli at each position on mesoglea were determined by averaging over the 8 measurements. Data points with standard deviation larger than 40 % were discarded. The relative measurement position was determined using ImageJ [61].

### Nano-indentation analysis

The bulk elastic modulus *E* was calculated with the modified Hertz model [26]:

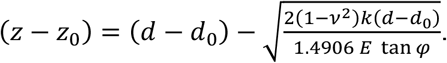

*z* is position of the tip, *d* cantilever deflection. *z_0_* and *d_0_* are the values corresponding to the first tip-sample contact. *v* is Poisson’s ratio (0.5), *k* the spring constant of the cantilever, and *φ* the half-opening angle of the AFM tip. To determine the elastic modulus, we have the contact point candidates “march” and applied the algorithm by Lin et al. [28] and Dimitriadis et al. [27]. The minimization of mean sum of square residuals (SSR) yielded the best-fit result (Figure 2).

### Mass spectrometry

Proteome samples were prepared as described in SI Material and Methods and analyzed in three biological replicates on an LTQ-Orbitrap XL mass spectrometer coupled to a nanoAcquity ultra performance LC system. The reverse phase-LC system consists of a 5 μm SymmetryC18 pre column and a 1.7 μm BEH130 C18 analytical column. Peptide mixtures were loaded on the pre column at a flow rate of 7 μL/min and were then eluted with a linear gradient at a flow rate of 0.4 μL/min. The mass spectrometer was operated in the data-dependent mode to measure MS and MS2 automatically. LTQ-Orbitrap XL was set to acquire a full scan at 60000 resolution at 400 m/z from 350 to 1500 m/z and simultaneously fragment the top 6 peptide ions in each cycle in the LTQ. The selected ions were excluded from MS/MS for 35 seconds.

## Acknowledgements

We thank ESRF for the synchrotron beamtime. M.V. thanks M. Sontag González and F. Gebert for experimental assistance. This work was supported by the German Science Foundation (DFG) through the Collaborative Research Centers CRC 1324 (A5 to T.W.H.), CRC 873 (A1 to T.W.H. and B7 to M.T.), the C.H.S. foundation (T.W.H. and H.P.), and the Germany’s Excellence Strategy –2082/1 – 390761711 (to M.T.). M.V. thanks DFG for fellowships (GRK1114 and EcTop2). M.T. thanks the JSPS (KAKENHI Grant Numbers 19H05719 and 20H00661) and Nakatani Foundation for supports.

## Author contributions

M.T. and T.W.H. designed and directed the research. M.V., S.K., and S.Ö, cultured polyps and prepared mesoglea samples. M.V., W.A., and M.B. performed non-GISAXS experiments and M.V. measured and analyzed AFM indentation. R.S. performed live imaging of *Hydra* polyps. M.V., T.W.H. and M.T. wrote the manuscript and all authors were involved in the discussion.

