## Supplemental Figures S1-S3 for "Wnt/β-Catenin Signaling Controls Spatio-Temporal Elasticity Patterns in Extracellular Matrix during *Hydra* Morphogenesis"

**Culture synchronization**. *Hydras* with buds at different developmental stages were collected from a culturing box, which was fed for at least 2 - 3 weeks on a daily base and transferred to 24-well-plates (1 *Hydra*/ well). The detachment of new buds from collected collected *Hydra* was checked twice a day. To obtain a synchronized culture the freshly detached buds were collected and transferred to new 24-well-plates (1 *Hydra*/ well). The day of detachment was defined as $t$ = 0 d.

**Mass mesoglea isolation.** Large numbers of mesogleas were isolated by a modification of the method used by Shostak et al [1]. In order to inhibit proteolytic degradation during isolation, Protease inhibitor cocktail tablets (cOpmlete Ultra tablets, mini, *EASYpack*, Roche, Switzerland) were used according to the procedure protocol provided by the supplier. Pellets of 90-150 frozen *Hydra* animals were transferred to protease inhibitor cocktail and allowed to thaw at room temperature. After thawing, *Hydra* animals were transferred to a glass test tube with 3 – 4 mL protease inhibitor cocktail, and cells were removed by successive pipetting using a Pasteur pipette. Subsequently 1 – 2 ml of 80 % sucrose solution were gently injected below the suspension to prevent the mesogleas from sticking to the glass bottom. The mixture was then centrifuged at 4 °C and 300 g for 5 min using a swing-out bucket rotor. After centrifugation mesogleas were collected from the sucrose-water interface. The procedure was repeated 7 – 10 times.

**Nano-GISAXS data analysis**. After background subtraction and deadtime correction satellite peak positions were determined using 2D-gaussian fit. The error was estimated by the half width at half maximum. The angle was determined from the dot product of reciprocal space vectors according to

$$\cos\gamma^* = \frac{\overrightarrow{a^*} \cdot \overrightarrow{b^*}}{|\overrightarrow{a^*}||\overrightarrow{b^*}|} \quad \text{(S1)}$$

where $a^*$ and $b^*$ represent the reciprocal lattice vectors with an angle of $\gamma^*$. The angle in real space, $\gamma$, was calculated as

$$\gamma = \pi - \gamma^* \quad \text{(S2)}$$

The real lattice vectors were determined using $\vec{a} \cdot \overrightarrow{a^*} = 2\pi$ and $\vec{b} \cdot \overrightarrow{b^*} = 2\pi$.

**Elasticity pattern classification.** Average elastic moduli of upper gastric region ($0.6 \leq d \leq 1.0$), and peduncle ($0.0 \leq d < 0.1$) were normalized by the elastic modulus of the budding region ($0.1 \leq d \leq 0.3$). The distribution of $E_{gastric}/E_{budding}$ and $E_{peduncle}/E_{budding}$ are shown in Figure S1. Each histogram could be fitted with a sum of two Gaussians. In case of $E_{gastric}/E_{budding}$ the two Gaussians intersected at $E_{gastric}/E_{budding} = 0.71$ (Figure S1a). This value was taken as the threshold to discrimination between Type A and Type B/C: elasticity maps with $E_{gastric}/E_{budding} \geq 0.71$ were considered as uniformly elastic, i.e. type A patterns. The remaining patterns were compared with regard to the elasticity of peduncle relative to budding region, i.e. $E_{peduncle}/E_{budding}$. In This case the intersection point of the fitted Gaussians found at $E_{peduncle}/E_{budding} = 0.85$ (Figure S1b) served as criterion for discrimination between pattern B, with $E_{peduncle}/E_{budding} \geq 0.85$, and pattern C with $E_{peduncle}/E_{budding} < 0.85$.

**Definition of marching range**. The onset point was chosen as the center of the marching range. This point was determined roughly by fitting linear functions to the outermost parts of the noncontact and contact region of a force curve given by following relations, respectively:

$$r_{noncontact} = \left\{(z,d) \in data \middle| \frac{(z_{min} + z_{max})}{2} < z\right\} \quad \text{(S3)}$$

$$r_{contact} = \left\{(z,d) \in data \middle| \frac{(d_{min} + d_{max})}{2} < d\right\} \quad \text{(S4)}$$

where *z* and *d* corresponds to the piezo and deflection of a datapoint, $z_{max}$ and $z_{min}$ corresponds to the maximum and minimum piezopositions and $d_{max}$ and $d_{min}$ represent the maximum and minimum cantilever deflection in the dataset. Subsequently, the data point nearest to the intersection of the two fitted lines was defined as the center of marching range ($z_c$, $d_c$). All datapoint within a specific radius *R* from the inflection point were assigned to the marching range.

$$s_{marching} = \left\{(z,d) \in data \middle| \sqrt{(z - z_c)^2 + (d - d_c)^2} \leq R\right\} \quad \text{(S5)}$$

The radius was adjusted according to the force curves so that the points at the beginning and end of the marching range were definitly far from the transition from contact to non-contact region.

**Alsterpaullone treatment.** Freshly detached buds collected from a synchronized culture were incubated in a 5 µM solution of alsterpaullone (Sigma-Aldrich, United States) in 0.025% dimethyl sulfoxide (Sigma-Aldrich, United States) in *Hydra* medium for 1 d. Subsequently, polyps were washed several times with *Hydra* medium and cultured for up to 4 d.

**Sample preparation for proteome analysis.** Isolated mesoglea from wild type and SILAC *Hydra* animals were precipitated by adding 4 × volumes of 80% acetone (– 20 °C) and stored over night at − 20°C. Samples were washed with 80 % acetone and the pellets were dissolved in 6 M guanidine HCl in 50mM TrisHCl (pH = 8.4). DTT was added to a final concentration of 10 mM, and the samples were incubated for 5 min at 95 °C. For the alkylation of reduced cysteins iodoacetamide was added to a final concentration of 20 mM and stored for 30 min at RT. To quench the remaining iodoacetamide, the addition of DTT to a final concentration of 10 mM was added. Samples were diluted by 50 mM Tris (pH = 8.4). Lys-C was added in an enzyme sample ratio of 1:50 and incubated at 25°C for 18 h. After digestion, samples were acidified with TFA and desalted with homemade SPE cartridges using POROS R2 material (Applied Biostystem). Peptides were eluted with 80% acetonitrile, dried in a speed vac and subsequently re-suspended in water. Peptide concentration were measured at 205 nm wavelength on a NanoDrop instrument and non-labeled and SILAC samples were mixed 1:1.

**Mass spectrometry data analysis.** LC-MS/MS raw data were analyzed with MaxQuant version 1.4.0.3 [2] with default settings. Proteins were identified using NCBI Hydra *magnipapillata* proteome database (Annotation release.101). Cysteine carbamidomethylation was used as a fixed modification and methionine oxidation, protein N-terminal acetylation as variable modifications. For the identification, the false discovery rate was set to 0.01 for peptides and proteins (the minimal peptide length allowed was six amino acids).

Protein ratios were normalized by its own “Spike-in” ratio as described in M. Looso *et al.*[3]. Using Perseus, protein ratios were additionally normalized for systematic errors by dividing with the median ratio. Protein IDs were filtered for proteins with at least 3 quantitative values in at least one time point.

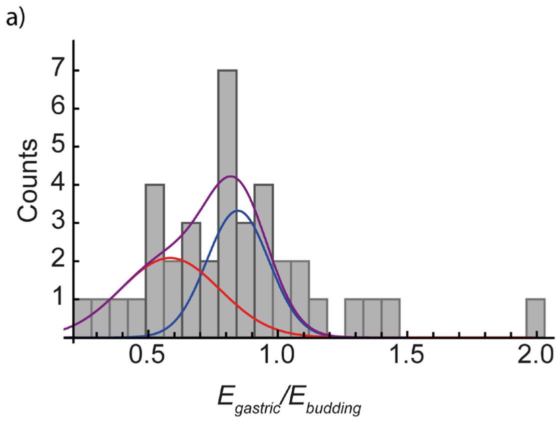


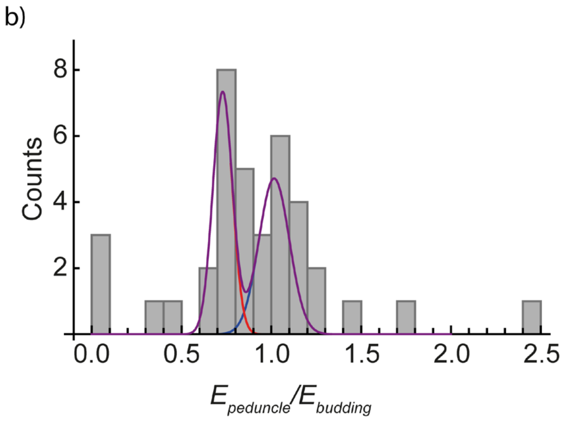


**Figure S1.** Distribution of elastic moduli of a) upper gastric region and b) peduncle normalized by the elastic modulus of budding region. A multi-peak Gaussian was fitted to each histogram (purple). Red and blue curves represent Gaussian functions contributing to the multi-peak fit. The red and blue curves intersect at (a) $E_{peduncle}/E_{budding}$ = 0.71 and (b) $E_{peduncle}/E_{budding}$ =0.85.

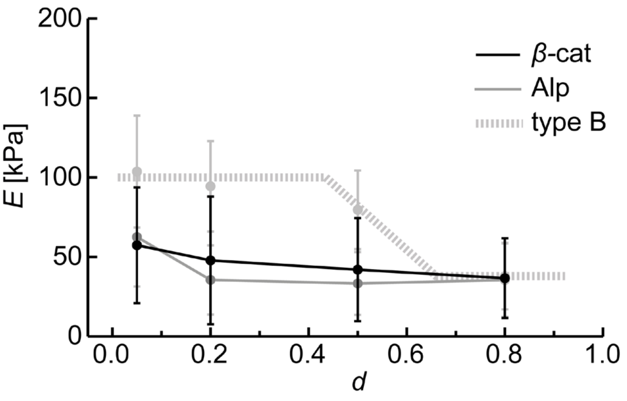


**Figure S2.** Average elastic modulus along the body column of β-catenin overexpressing Hydras (black). Although β-catenin overexpressing Hydras showed a broad variety of elasticity patterns in consent with a large phenotype diversity, the average elastic moduli ($n$ = 27) over 8 days compare well with those of Alp-type Hydras (grey). Type B elasticity pattern obtained for wild-type Hydras is shown by dashed gray line for comparison (see figure 3). Each data point represents the mean elastic modulus over the following regions: peduncle ($0 \leq d < 0.1$), budding region ($0.1 \leq d \leq 0.3$), center ($0.4 \leq d < 0.6$) and gastric region ($0.6 \leq d \leq 1.0$). Error bars correspond to standard deviations.

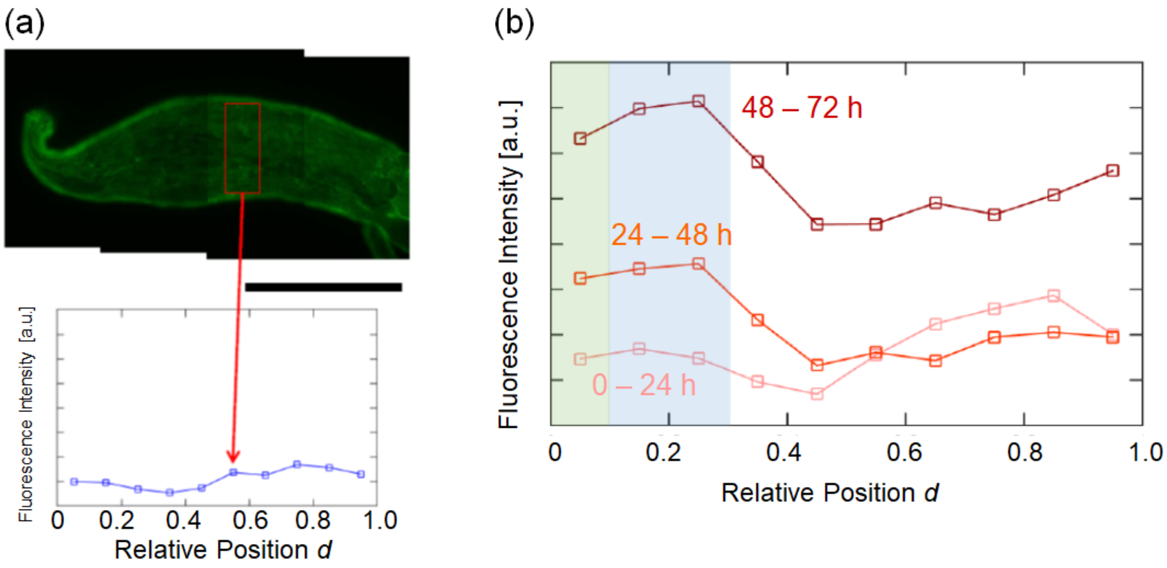


**Figure S3.** Distribution of LifeAct GFP signals along the body axis of LifeAct ecto *Hydra* at $t$ = 0 h. (a) Each data point represents the signal integrated inside the box. (b) Fluorescence intensities collected by live imaging every 6 h were averaged.

### References to Supporting Information


1. Shostak, S., Patel, N.G., and Burnett, A.L. (1965). The role of mesoglea in mass cell movement in Hydra. Developmental Biology *12*, 434-450.
2. Cox, J., and Mann, M. (2008). MaxQuant enables high peptide identification rates, individualized p.p.b.-range mass accuracies and proteome-wide protein quantification. Nat Biotech *26*, 1367-1372.
3. Looso, M., Michel, C.S., Konzer, A., Bruckskotten, M., Borchardt, T., Krüger, M., and Braun, T. (2012). Spiked-in Pulsed in Vivo Labeling Identifies a New Member of the CCN Family in Regenerating Newt Hearts. Journal of Proteome Research *11*, 4693-4704.
